# Analysis of miRNA inhibitors Efficacy using Capillary Electrophoresis with Laser-Induced Fluorescence

**DOI:** 10.1101/566786

**Authors:** Dong-Kyu Chae, Eunmi Ban

## Abstract

In the recent years, microRNAs (miRNAs) have been discovered to play a very important role in biological processes such as development, differentiation, and apoptosis. The miRNA expression levels in cells are associated with diverse diseases including cancers. MiRNA inhibitors have been widely employed for studying the functions and targets of miRNAs by transfecting the inhibitors into cells. The concentrations of miRNA inhibitors used for such studies can vary depending on the types of miRNAs being tested, the cell lines under study, and the analysis methods. Therefore, in order to obtain accurate results, appropriate amounts of miRNA inhibitors have to be used in the experiments. Apart from amounts, the evaluation of inhibitors may also have to be conducted for functional studies.

Here we developed capillary electrophoresis with laser-induced fluorescence (CE-LIF) method for evaluating miRNA inhibitor and for optimizing miRNA inhibitor concentrations, in the A549 lung cancer cell line. The target miRNAs, miRNA-23a and miRNA-24 are biomarker candidates in lung cancer cell lines. Our results showed that miRNA-23a and miRNA-24 were effectively inhibited upon transfection with 20 nM miRNA inhibitors using CE-LIF method. Furthermore, these results demonstrated the potential of CE for fast, specific, sensitive and specific analyses for the evaluation and determination of the optimal concentration of miRNA inhibitors for functional studies.

**Abstract Graphics:** 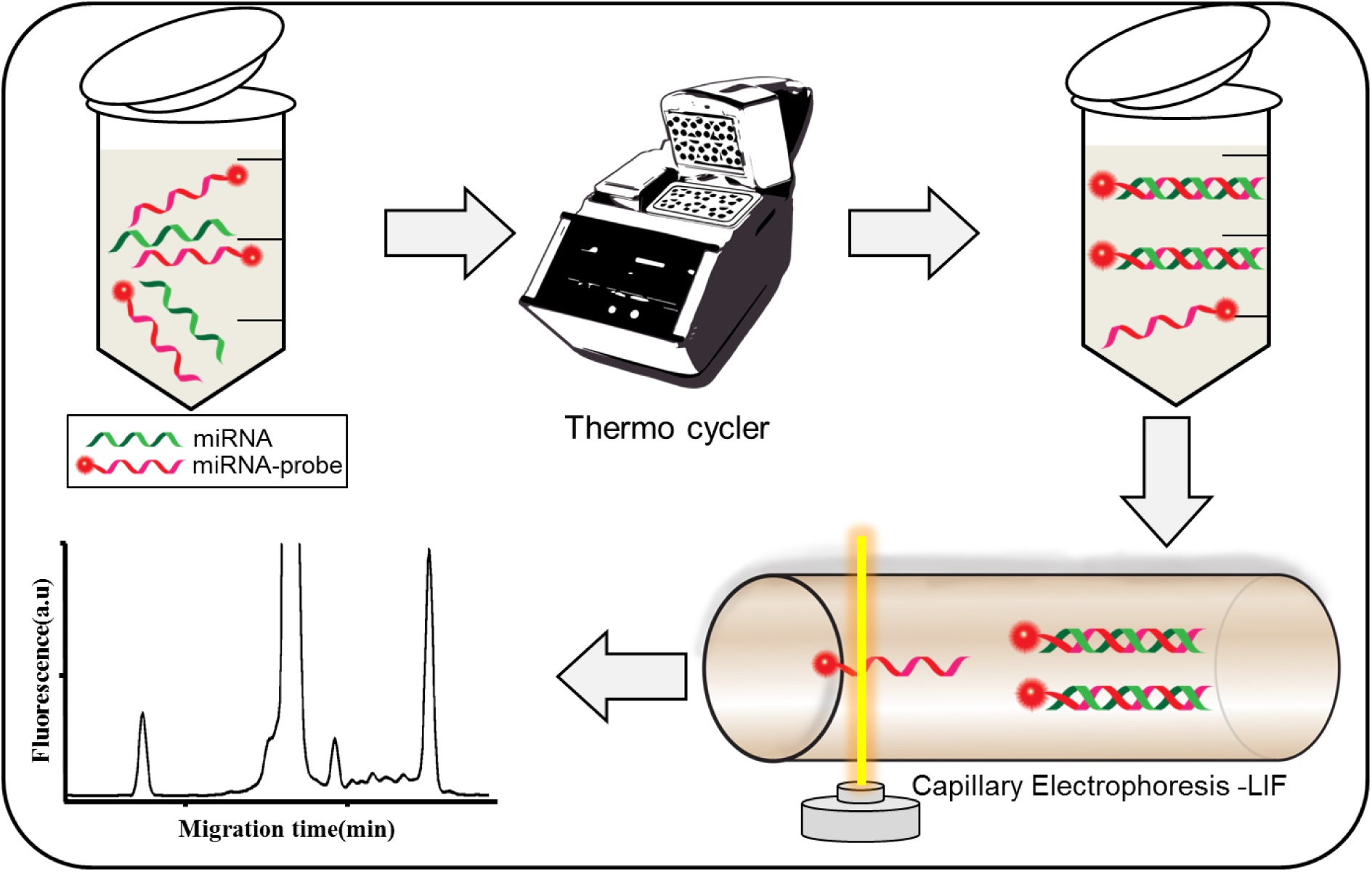

## 1. INTRODUCTION

MicroRNAs (miRNAs) are small, non-coding RNAs that are transcribed from genomic DNA into long primary transcripts (pri-miRNAs) by RNA polymerase II. Pri-miRNAs are recognized by the Drosha enzyme along with the DGCR8 protein and cleaved into the pri-miRNA stem-loop structure of the precursor miRNA (pre-miRNA) [1, 2]. Pre-miRNAs are transported from the nucleus to the cytoplasm by Exportin-5 and Ran-GTP, and are further processed by the Dicer-TRBP complex into mature miRNA duplexes [3, 4]. One strand of the miRNA duplex is loaded into the RNA-induced silencing complex (RISC), followed by base pairing with its target mRNA sequence in the 3’ UTR, resulting in the silencing of gene expression, either by inhibition of protein translation or degradation of the target mRNAs [5-8]. Consequently, miRNAs are associated with gene regulation, apoptosis, carcinogenesis, and metastatic diseases.[5, 9] Many studies have reported altered expression levels of miRNAs in various diseases, which shows the potential use of regulating miRNA levels in therapeutics. Therefore, a systematic functional study of miRNA regulation should be performed for the development of miRNA-based therapeutic drugs.

Loss-of-function strategies are widely used for studying miRNA function. There are several approaches for inhibiting miRNA function [10, 11], including gene knockouts in animals [12], miRNA sponge [13-15], and anti-miRNA oligonucleotides. MiRNA sponges have multiple miRNA-binding sites and simultaneously bind several miRNAs. In addition, the expression of miRNA sponges can be maintained over an extensive period, which results in a long-term inhibition in cells. MiRNA inhibitors are the most widely used, loss-of-function tools for studying miRNA functions. These inhibitors are transfected into cells where they bind target miRNAs that are loaded into Argonaute 2 proteins in the RISCs. The miRNA-induced silencing complex (miRISC) binds to the 3′ UTR regions of the target mRNAs [16-18], consequently regulating miRNA functions. MiRNA inhibitors should be stable in *vivo*, have high binding affinities to the target RNAs, and display a high but specific uptake in tissues for effective *in vivo* regulation. Commonly, miRNA inhibitors have been chemically modified by 2′-O-methylation and by using locked nucleic acids (LNA) oligonucleotides [19-21]. Recent functional studies involving miRNA inhibitors of the let-7 family have shown that the inhibitors mutually target miRNAs within the family, causing off-target effects; whereas, high doses of the inhibitors induce loss of specificity [22]. It has also been shown that inaccurate doses of the miRNA inhibitors are cytotoxic to the cells, or could induce a change in the cellular metabolism.[23, 24] Moreover, the performance of each miRNA inhibitor can vary depending on the type of targeted miRNA, the cell line being used, and the manufacturer [22, 25]. Therefore, for accurate functional studies of miRNA, it is necessary to optimize transfection conditions and evaluate inhibition efficiencies of the miRNA inhibitors.

The transfection conditions for miRNA inhibitors have been commonly optimized by recording the resultant phenotypic responses; however, such results are not accurate because cross-reactivity or specificity problems are not directly reflected in these observations. In addition, miRNA inhibitors are usually evaluated by northern blotting, quantitative real-time polymerase chain reaction (qRT-PCR), or luciferase assays. However, each method has its own limitations. Northern blotting displays low sensitivity and difficulty in separating the inhibitors from the heteroduplexes. Even though qRT-PCR is widely used for the evaluation of miRNA inhibition, the data from this technique could incorrectly indicate a higher inhibition of miRNA due to masking effects [16]. The limitations of the luciferase assay include the short half-life of the fluorescence and the difficulty in detecting the expression of more than two genes [26]. Therefore, new methods for optimizing transfection conditions and evaluating miRNA inhibitors need to be developed.

Capillary electrophoresis with laser-induced fluorescence (CE-LIF) has recently been used for the direct detection of miRNA levels in biological samples [27-30]. The advantages of this technique include superior resolution, quick analysis, and accurate quantification. The determination of miRNAs in biological samples by CE-LIF has been consistently reported in literature. Moreover, recent studies have shown that extremely low concentrations of miRNAs could also be detected by this method [27, 31]. Therefore, in this study we have used CE-LIF to evaluate miRNA inhibitors for use in the functional studies of miRNAs.

In this study, the CE-LIF method was developed for determining the optimal transfection conditions for functional studies. The expressions of miRNA-23a and miRNA-24 are known to be upregulated in lung cancer cells [24, 32]; therefore, they are useful biomarker candidates for lung cancer. After the transfection of cultured lung cancer cells with miRNA-23a and -24 inhibitors under optimized transfection conditions, cell viabilities were examined by performing the 3-(4,5-Dimethylthiazol-2-yl)-2,5-diphenyltetrazolium bromide (MTT) assay.

## 2. EXPERIMENTAL SECTION

### 2.1 Cell culture and RNA extraction

A549 and H358 cells were purchased from Korean Cell Line Bank (KCLB 10185, 25807). Cells were maintained in RPMI-1640 medium (Gibco, Carlsbad, CA, USA) supplemented with 10% FBS and 100 units/mL penicillin-streptomycin. The supplemented medium was changed every two days. Cells were grown under the standard incubation conditions of 37 °C, 5 % CO_2_, 95 % air, and 95% relative humidity. Cells were seeded in 100-mm petri dishes at a density of 1 × 106 cells/dish and were cultured for 1 day.

The day after seeding, cells were washed twice with cold Dulbecco’s PBS, scraped off from plates, and pelleted by centrifugation at 1400 ×g for 5 min at 4°C. The miRNA was extracted by using the Trizol reagent (Invitrogen, CA, USA) according to the manufacturer’s instructions. The extracted miRNAs were analyzed for RNA contents by A260/A280 nm reading using BioPhotometer (Eppendorf, Germany). DNase/RNase free water (Invitrogen, CA, USA) was used to prepare all buffer solutions and to adjust synthetic DNA and RNA concentration. The miRNA samples were stored at –80°C.

### 2.2 Oligonucleotide transfection

MiRNA-23a inhibitor, miRNA-24 inhibitor and control inhibitor were designed and synthesized by Genolution (Seoul, Korea). A549 cells were seeded in 60 mm plate 1 day before transfection. Inhibitors were transfected into cells using Lipofectamine-2000 (Invitrogen, CA, USA) according to the manufacturer’s instructions. Analysis was performed 1 – 9 days after transfection.

### 2.3 Hybridization condition

Fluorescence-labeled singe-stranded DNA (ssDNA) probes were purchased from Cosmogenetech (Seoul, Korea), and 3’-terminals of the probes were labeled with 5’-carboxyfluorescein phosphoramidite (6-FAM). Synthetic HPLC-grade miRNA and DNA oligonucleotides (listed in Table S1) were purchased from ST Pharm (Seoul, Korea).

For hybridization, 6-FAM-labeled DNA probes (1 nM) were mixed with synthetic miRNAs (1 nM) or total RNAs in the hybridization buffer (50 mM Tris–acetate, pH 8.0, containing 50 mM NaCl, 0.1 mM EDTA, and 1% Triton X-100). The samples were incubated in a thermal cycler (Eppendorf) for denaturation at 95°C for 5 min, followed by a renaturation step at 40 °C for 15 min prior to introduction into the capillary.

### 2.4 Capillary electrophoresis

CE experiments were performed on a PA 800 plus CE system (Beckman Coulter, Fullerton, CA, USA) equipped with a LIF detector. Fluorescence was detected by excitation at 488 nm using a 3-mW argon-ion laser (emission at 520 nm). Separations were performed using an untreated capillary (Beckman Coulter), which had a 75-μm inner diameter and 40-cm length (30-cm effective length). Separations were performed using a running buffer of 125 mM Tris– borate (pH 10.0) containing 2.5 M urea at 25 °C by applying 14 kV while the sample compartment was maintained at 25 °C. Samples were introduced hydrodynamically at 0.5 psi for 10 s.

### 2.5 MTT assay

Cell viability was measured by MTT assay. A549 cells were seeded in 96 well plates at 2000 cells per well and incubated for 1–5 days after transfection. A549 cells were treated with MTT (5 mg/mL, Sigma) for 4 h at 37 °C, and then dimethyl sulphoxide (DMSO) was added into each well for 30 min. The absorbance at 570 nm was measured using a SpectraMax M3 microplate reader (Molecular Devices, CA). The relative cell viability was calculated by comparison to the control cells. The values shown are the mean ± SD from triplicate experiments.

## 3. RESULTS

### 3.1 Expression of miRNA-23a and -24 in lung cancer cells detected using CE-LIF

In order to select a suitable lung cancer cell line for the functional studies of miRNA-23a and -24 as biomarker candidates for lung cancer diagnosis, the expression levels of miRNA-23a and -24 were first analyzed in two lung cancer cell lines with different degrees of differentiation and metastatic abilities. Lung cancer cell lines H358 (undifferentiated) [33] and A549 (high metastatic ability) [34, 35] were tested, because miRNA expression levels could vary in cells that had widely different characteristics. The expression levels of miRNA-23a and -24 in extracts from H358 and A549 cells were determined by CE-LIF. The extraction method and CE separation condition are adapted to that described in previously published our article, but with minor modifications [30]. Also, in our previous study, we demonstrated that our method was highly specific and linearity [30]. The detection of miRNA-23a and -24 in cell extracts was carried out by hybridizing fluorescent-labeled ssDNA probes to their miRNAs, as shown in Figure 1. After hybridization, the samples were applied to the capillary columns, and the hybridized complexes were separated from the unhybridized probes based on charge to mass ratio, at constant voltage. Figure S1 shows an electropherogram plotted by analyzing a hybridization mixture containing 1 nM of the 6-FAM-labeled ssDNA probes and the two miRNAs, miRNA-23a and -24. The separation proceeded rapidly, and the peaks representing the complexes and the ssDNA probes were well separated. The two miRNAs, miRNA-23a and observed that the expression levels of miRNA-23a and -24 were relatively higher in A549 cells -24 could be separated in CZE mode by the charge and mass differences of miRNA because the length of miRNA-23a and -24 are 21 and 22 nt, respectively although the miRNAs have similar length. In addition, the low electrophoretic mobility of the hybrid complex as compared to the probe alone was because of a smaller charge-to-mass ratio. From this analysis, we than in H358 cells (Figure S2, Supporting Information), suggesting that miRNA-23a and -24 expressions may be associated with the degree of lung cell differentiation and metastatic ability. Therefore, all the subsequent studies related to the evaluation and optimization of the transfection conditions of miRNA inhibitors were conducted in A549 cells, using the CE-LIF method for loss-of-function studies.

**Figure 1.**
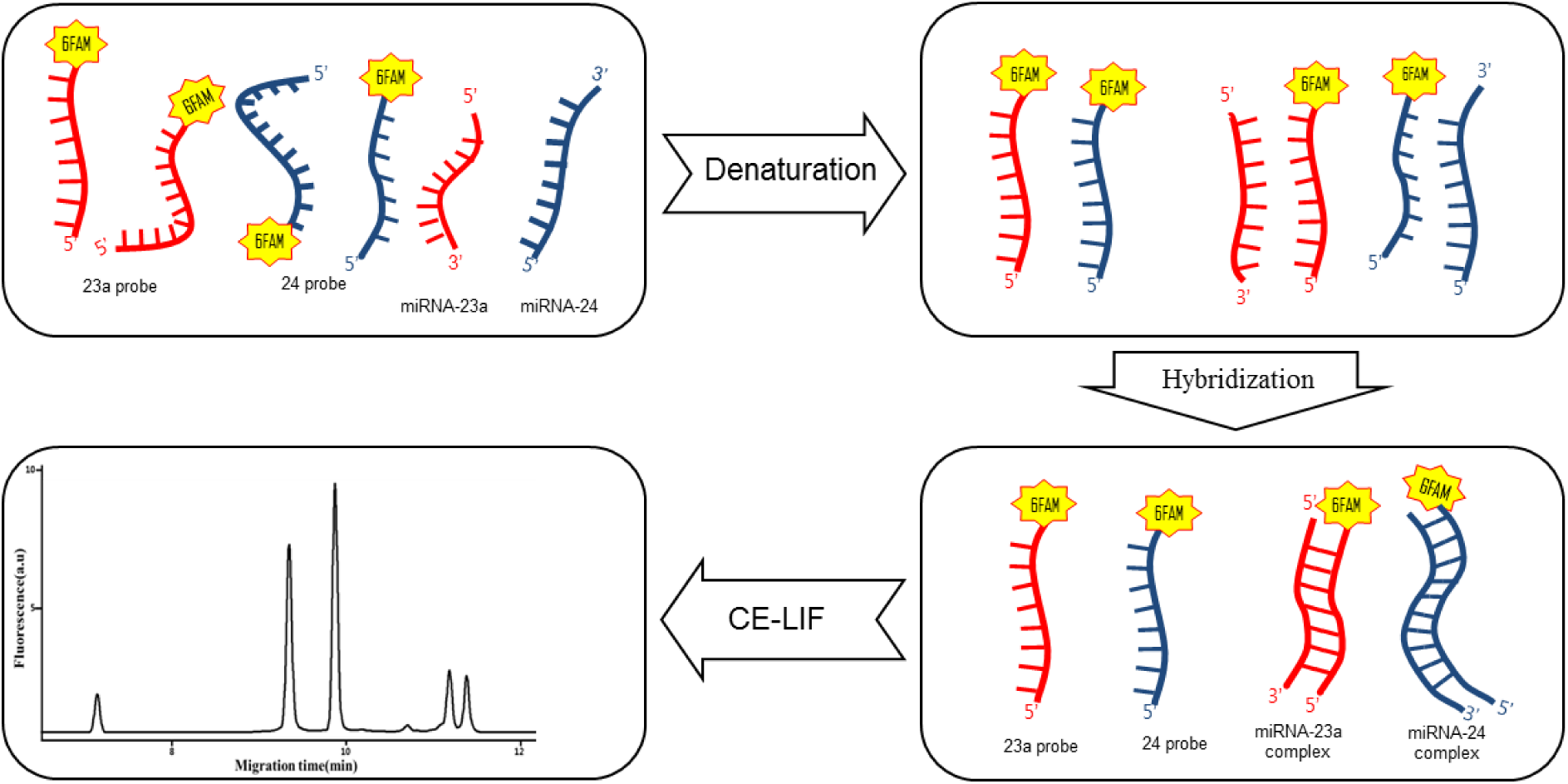
Schematic diagram for quantification of miRNA using CE-LIF.

### 3.2 Evaluation of miRNA inhibitors using CE-LIF

Before conducting miRNA loss-of-function studies using inhibitors, it is necessary to evaluate the ability of those inhibitors containing the target and control miRNA sequences to cause inhibition. In this study, the performances of miRNA-23a, -24, and control inhibitors were evaluated through the direct detection of endogenous miRNAs levels by using CE-LIF. The inhibitory properties of these inhibitors were confirmed by measuring peak intensities of the hybridization complexes of miRNA-23a and -24 with fluorescently labeled ssDNA probes in extracts from cells after transfection with miRNA-23a, -24, or control inhibitors. Results showed that in the control sample, transfection of cells with 20 nM control inhibitor resulted in the appearance of two peaks corresponding to endogenous miRNA-23a and -24. On the other hand, miRNA-23a levels in cells transfected with 20 nM miRNA-23a inhibitor decreased, but the miRNA-24 peak remained comparable to the peak in the control sample. Similarly, the levels of miRNA-24 also decreased by transfection with 20 nM miRNA-24 inhibitor, but the miRNA-23a peak remained almost unchanged (Figure 2).

**Figure 2.**
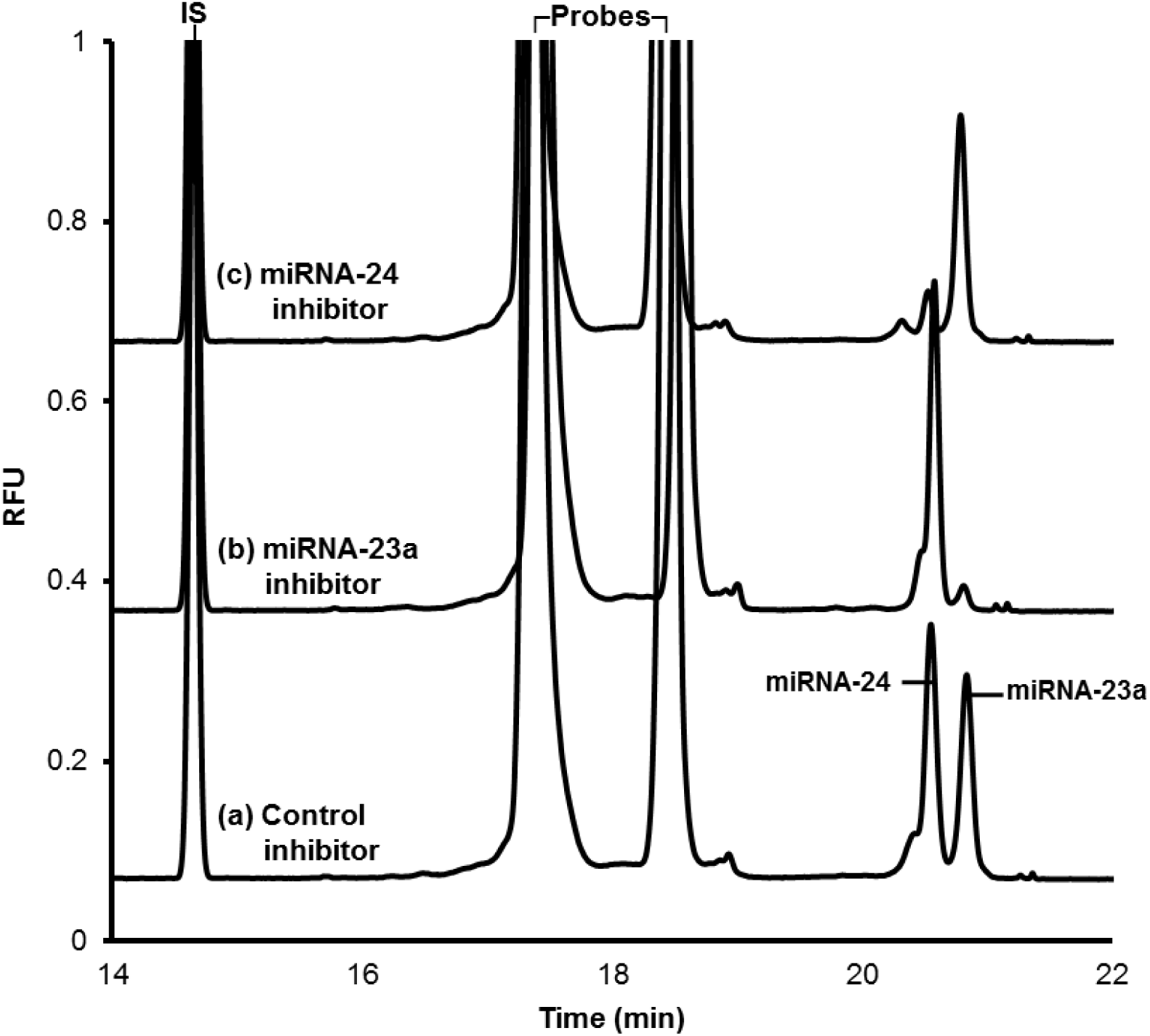
Electropherograms of miRNA-23a and -24. The total RNA from cell lysate was extracted and hybridized in hybridization buffer containing 1 nM miRNA-23a and -24 ssDNA probes after A549 cells were transfected with 20 nM of **(a)** control inhibitor, **(b)** miRNA-23a inhibitor, and **(c)** miRNA-24 inhibitor. Other conditions were same as in Figure S1.

From this electropherogram, we confirmed that the targeted miRNAs are effectively inhibited through specific binding between the miRNAs and the miRNA inhibitors. We also observed that the degree of inhibition effected by the two miRNA inhibitors is different. Thus, these results showed that CE-LIF could be used for the direct measurement of changes in miRNA levels through the binding of the miRNA inhibitors and target miRNAs, in contrast to other studies that measure these changes indirectly.

### 3.3 Optimization of miRNA inhibitors concentration for cell viability

To ensure reliable results during functional studies, the transfection conditions should be optimized. In particular, precise concentrations for transfection of miRNA inhibitors have to be determined, in order to maximize specific inhibition, and minimize cross-reactivity to non-target miRNAs. In this study, the concentrations of the miRNA inhibitors were determined by measuring the peak intensities of miRNA-23a and -24 from extracts of cells transfected with 10 to 40 nM of either the miRNA-23a or -24 inhibitors. Results showed that the degree of inhibition with the miRNA-23a and -24 inhibitors increased with increasing concentrations (Figure 3A and 3B). However, cross-reactions between the non-target miRNAs and the inhibitors were observed when the inhibitor was transfected at the highest concentration of 40 nM. These results imply that at very high concentrations, the miRNA inhibitors bind non-specifically to other miRNAs. Therefore, 20 nM was established as the optimal concentration for the transfection of miRNA inhibitors, in order to achieve the maximum inhibition possible with the least cross-reactivity.

**Figure 3.**
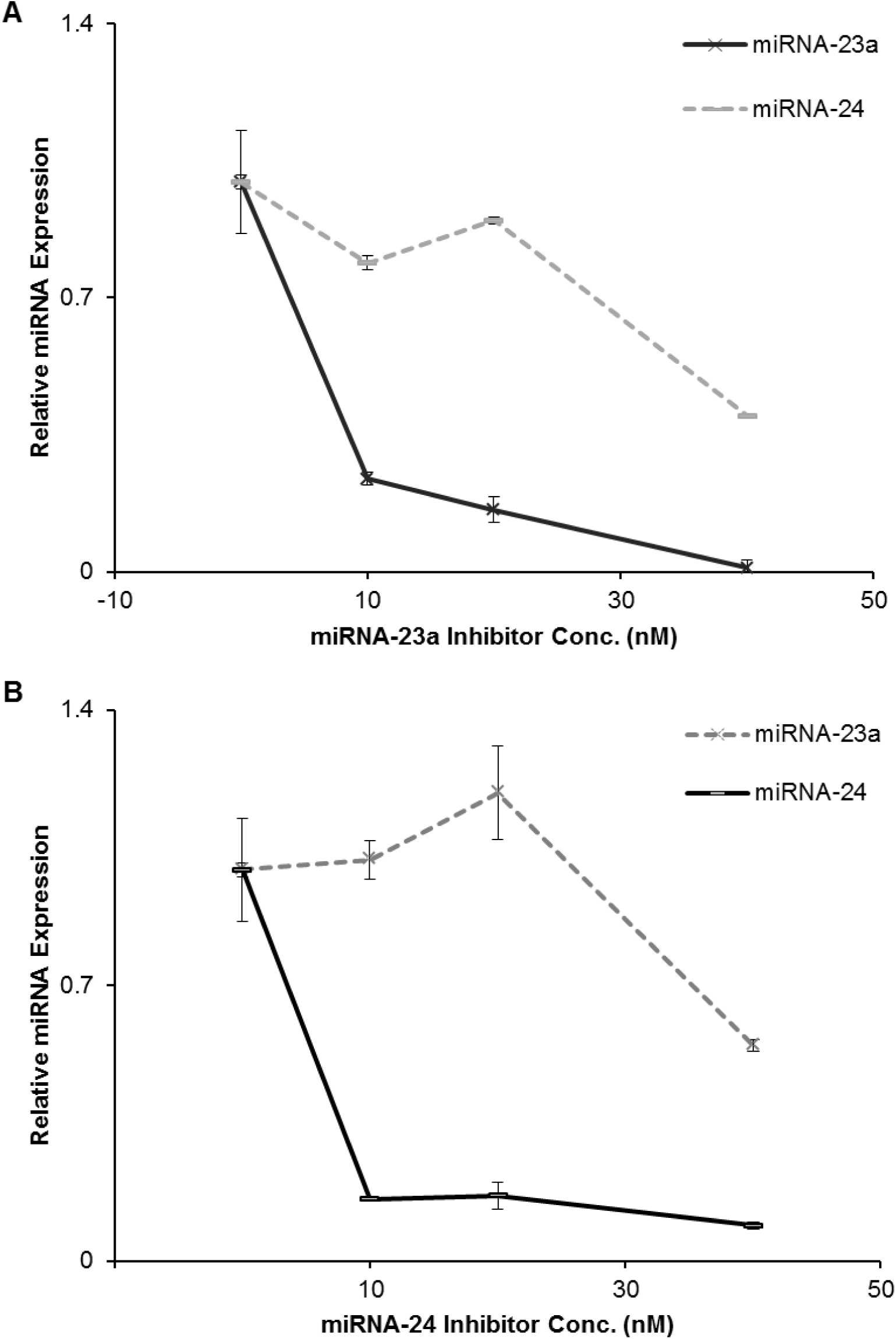
Levels of miRNA-23a and -24 expression. Total RNA were extracted from A540 cells after transfection **(A)** with miRNA-23a inhibitor, or **(B)** with miRNA-24 inhibitor. The concentration of miRNA-23a and -24 inhibitors was in between 10 nM to 40 nM and endogenous miRNA-23a and miRNA-24 levels were simultaneously detected using CE-LIF. All data were normalized on the levels of miRNA-23a and -24 from A549 cells transfected with 20 nM control inhibitor. Error bars represent mean and the SD from triplicate experiments.

### 3.4. Effects of miRNA inhibitors on cell viabilities

Since the optimal concentration determined for the miRNA inhibitors was 20 nM, this was used for further testing the effects of miRNA-23a and -24 on cell viability. Using the MTT assay, we confirmed that 20 nM miRNA-23a and -24 inhibitors inhibited the miRNA-23a and miRNA-24, resulting in restrained proliferation of cells. Figure 4 shows that the viability of the cells transfected with 20 nM miRNA-23a and -24 inhibitors was significantly lower than that of the cells transfected with the control inhibitor. As expected, this result showed that miRNA inhibitors accurately inhibited target miRNAs, which resulted in the decrease of cell viabilities. This result implies that miRNA-23a and miRNA-24 induce the proliferation of lung cancer cell. Therefore, our developed method for evaluation of miRNA inhibitors using CE-LIF could be applied for study of proper cellular phenotype such as apoptosis, migration, and invasion.

**Figure 4.**
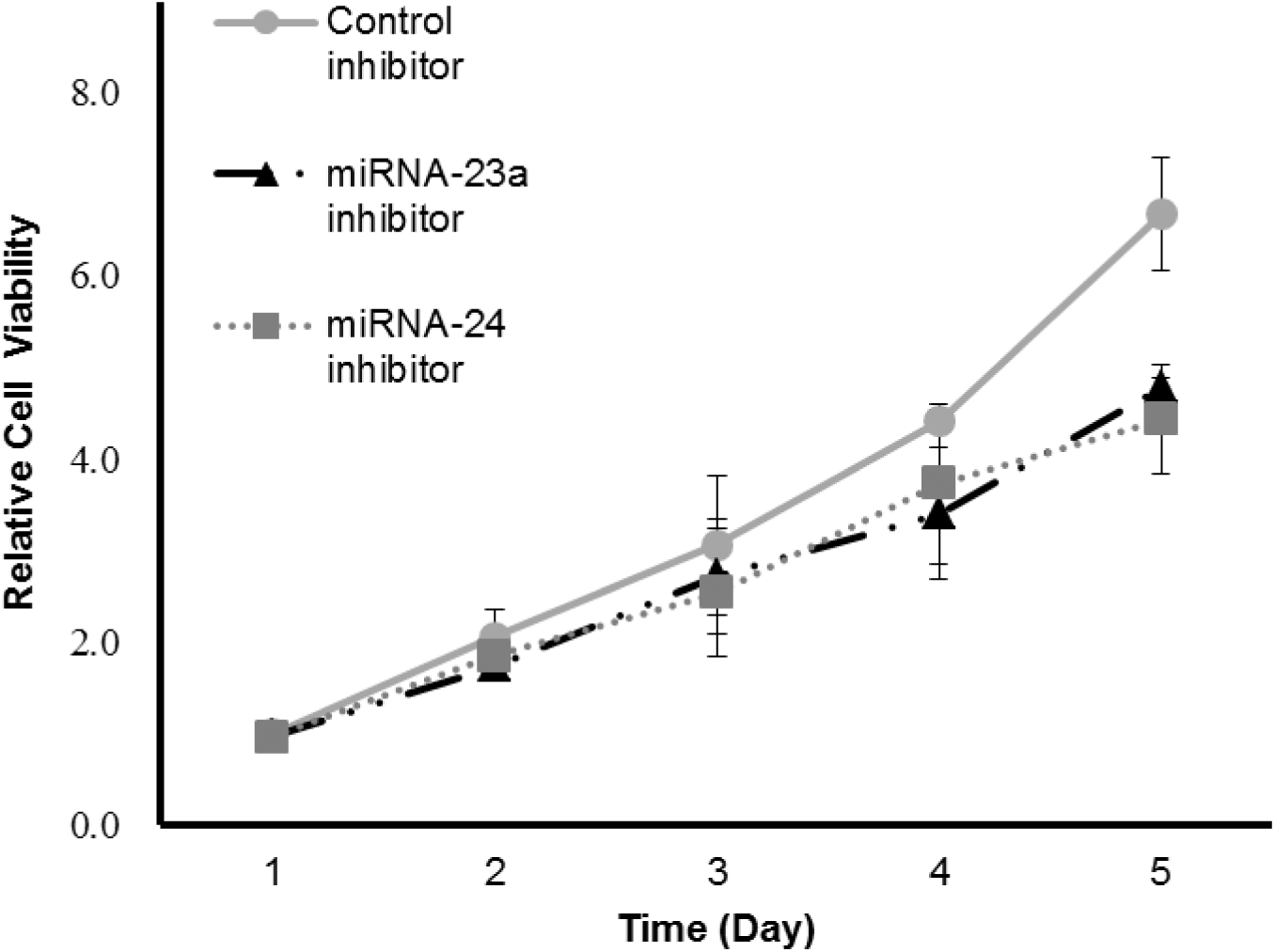
Cell viability of A549 cells. A549 cells were transfected with 20 nM miRNA-23a and -24 and control inhibitor, respectively. Cell viability was measured by the MTT assay 1–5 days after transfection. The relative cell viability was calculated by comparison to the control cells. Error bars represent mean and the SD from triplicate experiments.

## 4. DISCUSSION

The miRNA-23a and -24 genes exist in a cluster localized on chromosome 9q22, and they could be cooperating at the level of biological functions [36]. MiRNA-23a and -24 are expressed together but undergo post-transcriptional processing; therefore, the levels of both are different depending on the cell lines and biological conditions, and could be the cause of the cellular features of lung cancer cell lines [37]. Previous studies have reported that the miRNA-23a and -24 are simultaneously upregulated in various cancers, including lung cancer [24, 32]. The functions of miRNA-23a have been described in various diseases such as cancers, muscle atrophy, and cardiac disease [38-40]. In addition, miRNA-24 is also associated with the regulation of cell cycle and tumorigenesis [32]. Many studies have been performed on miRNA-23a and -24, but the functions of both continue to remain unclear. Generally, the study of miRNA function is conducted by transfection of miRNA inhibitors that have been modified in their backbone or sugar ring to reduce toxicity and improve affinity, specificity, and stability. It is possible, however that these diverse modifications may be contributing to the performance of the miRNA inhibitors. Moreover, the inaccurate use of these inhibitors might induce undesirable effects by targeting non-target miRNAs. Therefore, precise quantification and evaluation of the properties of these inhibitors should be carried out, in order to obtain accurate results for the study of miRNA function.

Figure 3 showed that high concentrations of the miRNA inhibitors could possibly cause cross-reactivity between them and non-target miRNAs. Usually, manufacturers of miRNA inhibitors and many other published studies have recommended that the inhibitors should be used at final concentrations between 1–50 nM, and warn that concentrations of >50 nM could be toxic to the cells. In addition, it is also advised that miRNA inhibitors should be used at optimized transfection conditions, i.e., optimal doses and treatment times should be used for miRNA functional studies [16, 41]. The optimization of transfection conditions are usually determined by observing phenotypic effects in cells, but phenotypes do not effectively represent the effects of cross-reactivity [20], e.g., it is difficult to conclude whether the phenotypic effects are due to the loss of non-target miRNAs. Thus, we evaluated the cross-reactivity of miRNA inhibitors by the direct detection of miRNAs, using CE-LIF. We confirmed that the miRNA-23a and -24 inhibitors inhibited the targets, miRNA-23a and -24, respectively, at concentrations ranging from 10–20 nM. However, at 40 nM, both miRNA-23a and -24 inhibitors showed cross-reactivity and inhibited both the miRNAs (Figure. 3).

Under optimal conditions, transfection of 20 nM miRNA-23a and -24 inhibitors in cells resulted in the reduction of cell viability, unlike that in case of the control inhibitor. Results from Figure 4 indicate that the miRNA-23a and -24 contribute to increasing cell viability.

## 5. CONCLUSION

In this study, CE-LIF method were adapted to determine the optimal transfection concentration of miRNA inhibitors for the functional study of miRNA-23a and -24 associated with lung cancer. Using CE-LIF, we observed that the miRNA-23a and -24 inhibitors inhibited the targets, miRNA-23a and -24, respectively, at concentrations ranging from 10–20 nM but at 40 nM, both miRNA-23a and -24 inhibitors showed cross-reactivity and inhibited both the miRNAs. The miRNAs regulate 60% of mammalian genes through binding to the 3’-UTRs and are associated with diverse diseases [42]. Recent studies show that miRNAs are important targets for therapeutic interventions. However, the development of miRNA therapeutics has challenges, such as miRNA delivery, undesirable interactions between the inhibitors and different targets. Therefore, direct measurements such as CE-LIF can contribute towards the accurate evaluation of the efficiency of miRNA inhibitors. In addition, the optimal concentrations of miRNA inhibitors should be determined by direct measurement methods such as CE-LIF, and not by observing the phenotypes. After all, it can be concluded that our study suggested a new evaluation method of miRNA inhibitor in order to obtain reliable results without unwanted interactions such as cross-reactivity for the functional studies of miRNA and the development of therapeutics using miRNA inhibitor.

## ACKNOWLEDGMENTS

We gratefully acknowledge the support of the GRL Program (20110021713) from the National Research Foundation of Korea and an institutional grant from Korea Institute of Science and Technology (2E24490 and 2E24860).

