## Supporting information for "Analysis of miRNA inhibitors Efficacy using Capillary Electrophoresis with Laser-Induced Fluorescence" (from another version of this preprint: version 3, 2019-03-15)

### Supplementary data

**Supplementary table S1** Overview of patient characteristics.

| Patient ID | Diagnosis | Histology | Stage (TNM) | Age at diagnosis |
|---|---|---|---|---|
| **1** | Ovarian cancer | Serous adenocarcinoma | pT2bNXM1 | 53 |
| **2** | Cervical cancer | Squamous cell carcinoma | pT4N0M0 | 74 |
| **3** | Cervical cancer | Endocervical adenocarcinoma | pT2bN0M0 | 56 |
| **4** | Uterine cancer | Uterine sarcoma | pT4N0M0 | 59 |
| **5** | Ovarian cancer | Serous adenocarcinoma | pT3aNXM0 | 75 |
| **6** | Cervical cancer | Squamous cell carcinoma | pT2bN1M0 | 48 |
| **7** | Ovarian cancer | Endometroid adenocarcinoma | pT1cN0M0 | 58 |
| **8** | Ovarian cancer | Serous adenocarcinoma | pT3cN1M1 | 65 |
| **9** | Ovarian cancer | Serous adenocarcinoma | cT3cN0M1 | 62 |
| **10** | Cervical cancer | Squamous cell carcinoma | pT4aN1M0 | 69 |
| **11** | Ovarian cancer | Clear cell adenocarcinoma | pT1aN0MX | 50 |
| **12** | Endometrial cancer | Endometrical adenocarcinoma | pT1aN0M0 | 57 |
| **13** | Ovarian cancer | Serous adenocarcinoma | pT3cN0M0 | 42 |
| **14** | Endometrial cancer | Endometrical adenocarcinoma | pT1aN0M0 | 46 |
| **15** | Vulvar cancer | Intraepithelial adenocarcinoma | cT3N3M1 | 54 |
| **16** | Uterine sarcoma | Uterine sarcoma | pT1bN0M1 | 73 |
| **17** | Uterine sarcoma | Carcinosarcoma | pT2aNXM0 | 55 |
| **18** | Endometrial cancer | Endometrical adenocarcinoma | pT1aN0M0 | 70 |
| **19** | Ovarian cancer | Microinvasive carcinoma | pT1aNXM0 | 91 |
| **20** | Ovarian cancer | Adult granulosa cell tumor | pT1aNXMX | 80 |

**Supplementary table S2** The 51 target genes in cancer panel.

| Gene name | Refseq | Total amount of Exon | Number of amplicons | Total amplicon length (bp) |
|---|---|---|---|---|
| ***ABL1*** | NM_007313 | 4/11 | 6 | 728 |
| ***AKT1*** | NM_001014431 | 13/14 | 25 | 2,908 |
| ***ALK*** | NM_004304 | 29/29 | 64 | 7,093 |
| ***AR*** | NM_000044 | 8/8 | 29 | 3,293 |
| ***BAP1*** | NM_004656 | 17/17 | 35 | 3,904 |
| ***BRAF*** | NM_004333 | 2/18 | 2 | 245 |
| ***BRCA1*** | NM_007300 | 23/24 | 73 | 7,779 |
| ***BRCA2*** | NM_000059 | 26/27 | 121 | 12,515 |
| ***BTK*** | NM_000061 | 1/19 | 1 | 127 |
| ***CCND1*** | NM_053056 | 5/5 | 13 | 1,470 |
| ***CD274*** | NM_014143 | 6/7 | 15 | 1,643 |
| ***CDK4*** | NM_000075 | 7/8 | 13 | 1,469 |
| ***CDK6*** | NM_001145306 | 7/8 | 16 | 1,775 |
| ***CDKN2A*** | NM_000077 | 2/3 | 8 | 935 |
| ***DDR2*** | NM_006182 | 16/18 | 74 | 8,679 |
| ***EGFR*** | NM_005228 | 7/28 | 9 | 1,112 |
| ***ERBB2*** | NM_004448 | 25/27 | 54 | 5,978 |
| ***ERBB3*** | NM_001982 | 5/28 | 6 | 742 |
| ***ERBB4*** | NM_005235 | 28/28 | 63 | 6,562 |
| ***ESR1*** | NM_001122740 | 8/9 | 27 | 3,149 |
| ***FGFR1*** | NM_001174067 | 18/19 | 39 | 4,364 |
| ***FGFR2*** | NM_022970 | 17/18 | 45 | 4,742 |
| ***FGFR3*** | NM_001163213 | 16/18 | 30 | 3,328 |
| ***FGFR4*** | NM_213647 | 16/18 | 30 | 3,548 |
| ***FLT3*** | NM_004119 | 4/24 | 4 | 485 |
| ***GNA11*** | NM_002067 | 4/7 | 10 | 1,163 |
| ***GNAQ*** | NM_002072 | 4/7 | 7 | 813 |
| ***HRAS*** | NM_001130442 | 2/5 | 2 | 187 |
| ***IDH1*** | NM_005896 | 1/10 | 1 | 123 |

| | | | | |
|---|---|---|---|---|
| ***IDH2*** | NM_002168 | 1/10 | 2 | 256 |
| ***KIT*** | NM_000222 | 9/21 | 10 | 1,217 |
| ***KRAS*** | NM_033360 | 3/6 | 3 | 336 |
| ***MAP2K1*** | NM_002755 | 2/11 | 4 | 485 |
| ***MET*** | NM_001127500 | 5/21 | 7 | 851 |
| ***MTOR*** | NM_004958 | 4/58 | 5 | 613 |
| ***MYCN*** | NM_005378 | 2/3 | 8 | 824 |
| ***NOTCH1*** | NM_017617 | 33/34 | 91 | 10,356 |
| ***NRAS*** | NM_002524 | 3/7 | 3 | 361 |
| ***PDGFRA*** | NM_006206 | 4/23 | 4 | 507 |
| ***PIK3CA*** | NM_006218 | 8/21 | 11 | 1,227 |
| ***PPARG*** | NM_015869 | 7/7 | 21 | 2,320 |
| ***PTCH1*** | NM_000264 | 23/24 | 59 | 6,493 |
| ***PTEN*** | NM_000314 | 9/9 | 17 | 1,753 |
| ***RAF1*** | NM_002880 | 4/17 | 4 | 500 |
| ***RET*** | NM_020975 | 7/20 | 57 | 6,638 |
| ***ROS1*** | NM_002944 | 7/43 | 7 | 854 |
| ***SMO*** | NM_005631 | 5/12 | 5 | 637 |
| ***STK11*** | NM_000455 | 9/10 | 16 | 1,924 |
| ***TP53*** | NM_001276760 | 8/11 | 22 | 2,478 |
| ***TSC1*** | NM_000368 | 21/23 | 49 | 5,521 |
| ***TSC2*** | NM_000548 | 40/42 | 87 | 9,826 |

**Supplementary figure S1.**

**A**

Variants in cfDNA

| Genes | Coding | Amino Acid Change | Ovarian | | | | | | | | | Cervical | | | | Endometrial | | | Uterine | | | Vulvar |
|---|---|---|---|---|---|---|---|---|---|---|---|---|---|---|---|---|---|---|---|---|---|---|
| *ALK* | c.2158C>G | p.L720V | | | | | | | | | | ■ | | | | | | | | | | |
| *AR* | c.173A>T | p.Q58L | ■ | | | | | | | | | | | | | | | | | | | |
| *BAP1* | c.1168C>A | p.P390T | | | | | | | | | | | | | | | ■ | | | | | |
| *BRCA1* | c.3548A>G | p.K1183R | ■ | | | | | | | | | | | | ■ | | | | | | | |
| | c.444G>T | p.Q148H | | | | | | | | | | | | | | ■ | | | | | | |
| | c.5080_5082delCAC | p.H1694del | | | | ■ | | | | | | | | | | | | | | | | |
| | c.1716_1717insA | p.S573fs | ■ | | | | | | | | | | | | | | | | | | | |
| *BRCA2* | c.5047C>T | p.Q1683* | | ■ | | | | | | | | | | | | | | | | | | |
| | c.3299A>C | p.N1100T | | | | | | ■ | | | | | | | | | | | | | | |
| | c.9364G>A | p.A3122T | | | | | | ■ | | | | | | | | | | | | | | |
| | c.3646T>G | p.F1216V | | | | | | | | | | | | | | | ■ | | | | | |
| | c.4334A>C | p.K1445T | | | | | | | | | | | | | | ■ | | | | | | |
| | c.7088A>G | p.Y2363C | | | | ■ | | | | | | | | | | | | | | | | |
| *BTK* | c.1475G>A | p.R492H | | | | ■ | | | | | | | | | | | | | | | | |
| *CDK4* | c.134G>C | p.G45A | | | | | | ■ | | | | | | | | | | | | | | |
| *ESR1* | c.437C>A | p.P146Q | ■ | | | | | | | | | | | | ■ | | | | | | | |
| *FGFR1* | c.947C>T | p.P316L | | | | | | | | | | | | | | | | | | | | ■ |
| *FGFR2* | c.238G>A | p.G80R | ■ | | | | | | | | | | | | | | | | | | | |
| *FGFR3* | c.1295G>A | p.S432N | | ■ | ■ | | | | | | | | | | | | | | | ■ | | |
| *MYCN* | c.134_135delCG | p.P45fs | | | | | | | | | | | | | | | ■ | | | | | |
| *NOTCH1* | c.2982C>G | p.N994K | | ■ | | | | | | | | | | | | | | | | | | |
| | c.6788G>A | p.R2263Q | | | | | | | | | | | | | | | ■ | | | | | |
| *PIK3CA* | c.3136G>A | p.A1046T | | | | | | | | | | | | | ■ | | | | | | | |
| *PTCH1* | c.3232C>T | p.L1078F | | | | | | | | | | | | | | ■ | | | | | | |
| *TP53* | c.743G>A | p.R248Q | | | | | ■ | | | | | | | | | | | | | | | |
| | c.785G>T | p.G262V | | | | ■ | | | | | | | | | | | | | | | | |
| | c.818G>C | p.R273P | | | | | | | | | | | | | ■ | | | | | | | |
| | c.602delT | p.L201fs | | | | | | | | | | | | | ■ | | | | | | | |
| | c.827C>A | p.A276D | | | | | | | ■ | | | | | | | | | | | | | |
| | c.722C>A | p.S241Y | | | | | | | | | | | | | | | | ■ | | | | |
| *TSC2* | c.2153G>A | p.R718H | | | | | | | | ■ | | | | | | | | | | | | |

**B**

Variants in ctcDNA

| Genes | Coding | Amino Acid Change | Ovarian | | | | | | | | | Cervical | | | | Endometrial | | | Uterine | | | Vulvar |
|---|---|---|---|---|---|---|---|---|---|---|---|---|---|---|---|---|---|---|---|---|---|---|
| *BAP1* | c.548T>C | p.L183P | ■ | | | | | | | | | | | | | | | | | | | |
| *BRCA1* | c.3113A>G | p.E1038G | | | | | ■ | | | | | | | | | | | | | | | |
| | c.329A>G | p.K110R | | | | | | | | | | | | | ■ | | | | | | | |
| | c.5207G>T | p.S1736I | | | | | | | | | | | | | | ■ | | | | | | |
| | c.2566T>C | p.Y856H | | | | | | | | | | | | | | | | | | | | ■ |
| | c.3114delA | p.A1039fs | | | | | | | | | | | | | | | | | | | ■ | |
| *BRCA2* | c.865A>C | p.N289H | ■ | | | | | | | | | | | | | | | | | | | |
| *CDK4* | c.753_754insC | p.R252fs | | | | | ■ | | | | | | | | | | | | | | | |
| *ERBB2* | c.1966T>C | p.S656P | | ■ | | | | | | | | | | | | | | | | | | |
| | c.1147G>A | p.G383R | | | | | ■ | | | | | | | | | | | | | | | |
| | c.1642C>T | p.Q548* | | | | | | | | | | | | | | | ■ | | | | | |
| *ERBB3* | c.1959_1960insT | p.V654fs | | | | | | | | | | | | | | | | | | | ■ | |
| *FGFR1* | c.1741delG | p.A581fs | | | | | | | | | | ■ | | | | | | | | | | |
| *FGFR3* | c.1900G>A | p.A634T | | | | | | | | ■ | | | | | | | | | | | | |
| *NOTCH1* | c.6551A>T | p.Q2184L | | | | | | | | | | | | | | ■ | | | | | | |
| *RET* | c.2308C>T | p.R770* | | | | | | | | | ■ | | | | | | | | | | | |
| *TP53* | c.31G>C | p.E11Q | | | | | | ■ | | | | | | | | | | | | | | |
| | c.214C>G | p.P72A | | ■ | | | | | | | | ■ | | | | ■ | | | ■ | | | |
| *TSC1* | c.1585G>A | p.A529T | | | | | | | | | | | | | | | | | ■ | | | |
| *TSC2* | c.2509G>A | p.A837T | | | | | | | | | | | | | | | | | | | | ■ |
| | c.3178T>C | p.W1060R | | | | | | | | | | | | | | | | | | | | ■ |
| | c.3299T>C | p.V1100A | | | | | | | | | | | | | | | | | | | ■ | |

**C**

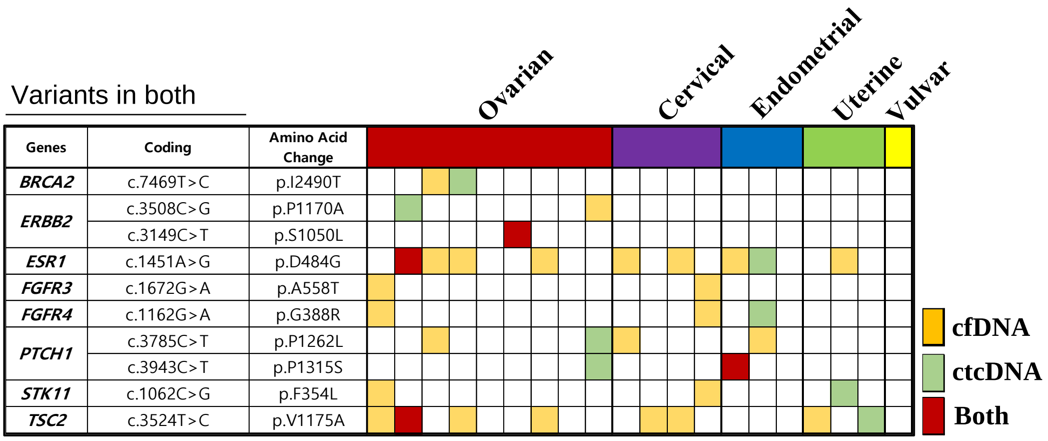


**Supplementary Figure S1** The analysis of genetic mutation for 20 patients for whom both cfDNA and CTCs samples were sequenced. **Note:** (A) Heat map showing the distribution of detected variants in cfDNA samples for each patient with the five main types of gynecologic cancer; ovarian, cervical, uterine, endometrial, and vulvar cancer. (B) Heat map representing variants in CTCs samples. (C) Heat map representing the overlapping variants detected in cfDNA and CTCs samples. Detected genetic variants in cfDNA (yellow), CTCs (green), and both (red).
